# Exploring the Frame Effect

**DOI:** 10.1101/2022.02.10.479960

**Authors:** Patrick Cavanagh, Stuart Anstis, Matteo Lisi, Mark Wexler, Marvin Maechler, Marius ’t Hart, Mohammad Shams-Ahmar, Sharif Saleki

**Affiliations:** Department of Psychology, Glendon College, Toronto, ON, Canada; Centre for Vision Research, York University, Toronto, ON, Canada; Department of Psychology, University of California at San Diego, La Jolla, CA, USA; Department of Psychology, Royal Holloway, University of London, London, UK; INCC and CNRS, Université de Paris, Paris, France; Department of Psychological and Brain Sciences, Dartmouth College, Hanover, NH, USA

## Abstract

Probes flashed within a moving frame are dramatically displaced (Özkan et al, 2021; Wong &Mack, 1981). The effect is much larger than that seen on static or moving probes (induced motion, Duncker, 1929; Wallach et al, 1978). These flashed probes are often perceived with the separation they have in frame coordinates — a 100% effect. Here we explore this frame effect on flashed tests with several versions of the standard stimulus. We find that the frame effect holds for smoothly or abruptly displacing frames, even when the frame changed shape or orientation between the endpoints of its travel. The path could be non-linear, even circular. The effect was driven by perceived not physical motion. When there were competing overlapping frames, the effect was determined by which frame was attended. There were a number of constraints that limited the effect. A static anchor near the flashes suppressed the effect but an extended static texture did not. If the probes were continuous rather than flashed, the effect was abolished. The observational reports of 30 online participants suggest that the frame effect is robust to many variations in its shape and path and leads to a perception of flashed tests in their locations relative to the frame as if the frame were stationary. Our results highlight the role of frame continuity and of the grouping of the flashes with the frame in generating the frame effect.

## Introduction

Vision has been shown to encode the motions and positions of objects relative to the frames that surround them (Duncker, 1929; Johansson, 1950). Frames can change what we judge to be “up” (Asch &Witkin, 1948; Morgan, Grant, Melmoth &Solomon, 2015) and what direction we think is straight ahead (Roelofs, 1935; Matin &Fox, 1989). A moving frame can alter the sense of our own motion (Warren, 1895) or that of an object within the frame (Duncker, 1929; Wallach 1959; Johansson, 1950). However, this earlier literature on frame effects typically examined static (Duncker, 1929) or continuously moving probes (Wallach et al, 1978). In contrast, moving frames give far larger effects for flashed rather than continuous probes (Wong &Mack, 1981; Özkan et al., 2021). The illusory offsets can be as large as the frame’s displacement, as if the flashes were seen in the frame’s coordinates and the frame were not moving. Moreover, unlike continuous probes, flashed probes do not appear to move. Johansson (1950) had proposed a process of motion decomposition for groups of elements in motion. This decomposition would extract a ‘‘common motion’’ vector shared by all the elements and leave the differences from the group vector as ‘‘relative motion’’. But flashed tests do not appear to have either the common or relative motion. Instead of a motion decomposition, we get a position decomposition and the effects are dramatic. This perception of the flash locations relative to the stabilized frame, rather than to their actual screen coordinates, may be linked to visual stability where the displacement of the entire visual scene acts as a moving frame that stabilizes position as the eyes move.

In this paper, we explore several variations of the effect of moving frames on flashed probes. Observational reports were collected from 30 online participants for 21 versions of the stimulus that varied the nature of the frame, the properties of its motion, and the grouping of the flashes with the frame. In most cases, the results are readily visible in the 21 movies. The observations we collected were simple subjective reports (e.g., yes, I see it) rather than controlled parametric tests, but they help identify which stimuli give strong effects and which give weak, ambiguous, or no effects. These results will then guide the selection of more controlled, future testing.

## Methods

### Participants

The experiment was conducted online. 30 participants were recruited by email sent to vision labs across the world. All 8 co-authors also participated. The protocols for the study were approved by the York University Review Board in accordance with the principles of the Declaration of Helsinki (2003).

### Stimuli and procedure

The experiment consisted of a set of 21 movies presented to the participants in their web browser, accessed here https://cavlab.net/Demos/FrameExperiment. The participants saw the same 21 videos of frame effects presented in the following results sections, each with a moving frame or background and two flashes, one red and one blue. The flashes were vertically aligned in most videos, separated center-to-center by approximately twice the diameter of the flashed discs. This vertical offset combined with any illusory horizontal offset served to create a noticeable angle between the upper and lower flashes that was more easily detected than the pure horizontal offset that would be generated by the frame effect for superimposed flashes. The offset was horizontal in one video that had vertical frame motion and in two other videos that had both horizontal and vertical motion components, the offset was oblique. The question for each test was presented below the video and there was no time limit. For most of the videos, the participants were asked whether the red flashed disc was seen to the right of the blue and they responded with a 4-point scale (1. Yes; 2. Yes, after a while; 3. Not much; and 4. No). Other responses were specific to different questions on some of the videos. They returned their responses by email. The experiment took 10 to 15 minutes.

### Analysis

The responses of the participants are combined and reported with the question and response choices for each of the 21 movies.

## Results

### Where Frames Work

#### 1.1.1. The basic frame effect

Here is the basic frame effect with an outline square moving left and right and a probe disc flashing at each reversal. Click the movie to run it in a browser window. The red and blue discs here are *always* vertically aligned — notice the green line — but when the line disappears, a large offset may appear with red to the right of blue by about as much as the frame’s displacement. All observers reported this offset.

*Do not stare at or fixate the flashes in any of the demonstrations, that will reduce or eliminate the effect*.

**Movie 1.1.1.**
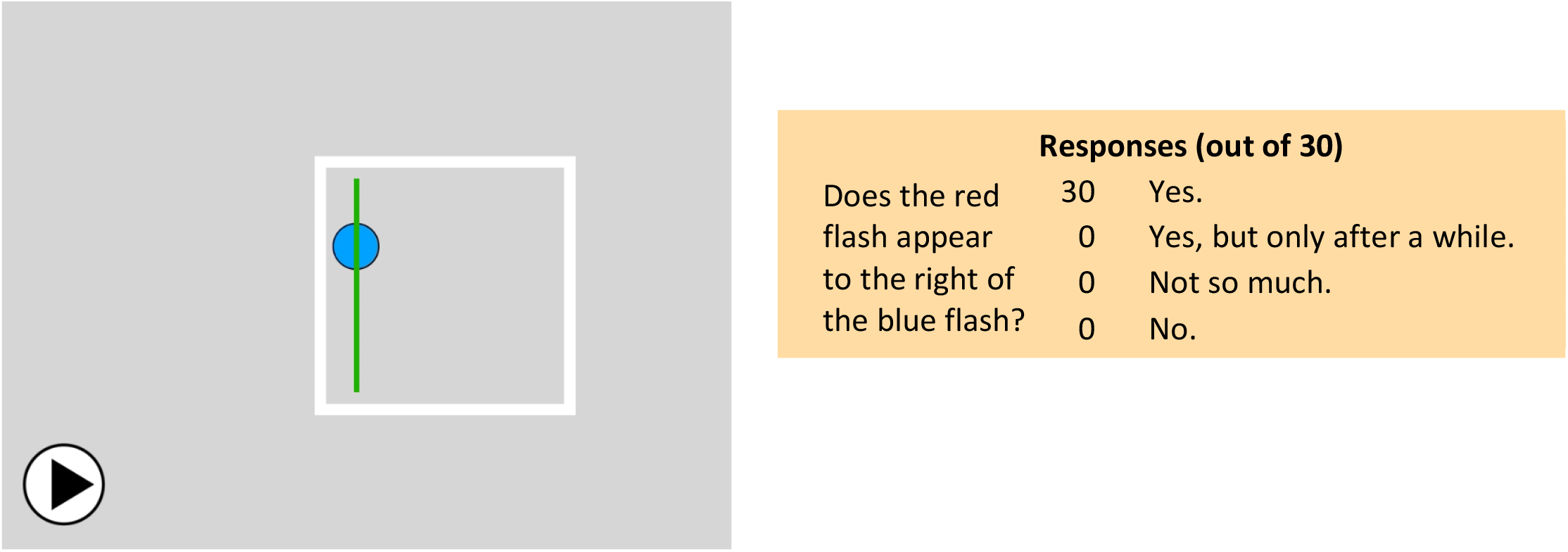
Click to start.

#### 1.1.2. Apparent motion

Here is the same basic frame effect but now with a single step in the displacement — apparent motion. The red and blue discs here are again *always* vertically aligned — notice the green line — but when the line disappears, the offset may appear with red to the right of blue. 76% of observers reported seeing this offset immediately or after some delay. 17% saw no offset.

**Movie 1.1.2.**
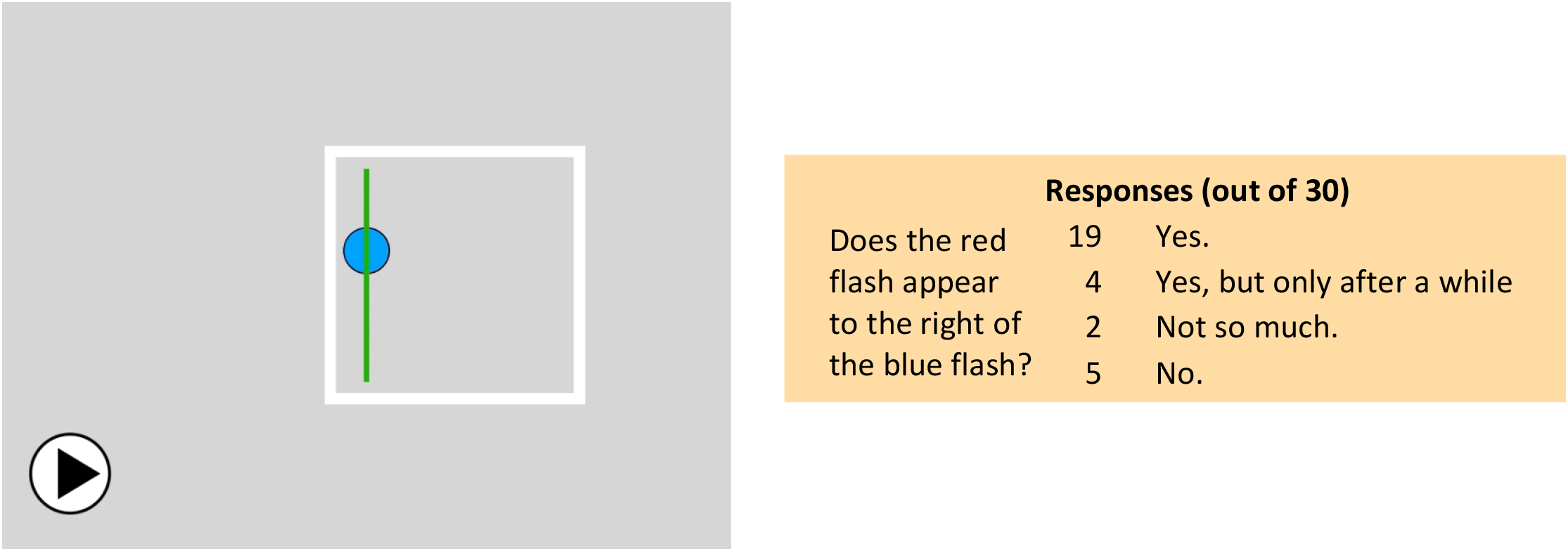
Click to start.

#### 1.1.3. Second-order motion

Here the frame is a second-order shape defined by flickering dots against the surrounding steady random dots (Cavanagh &Mather, 1989), there is no difference in mean luminance between the frame and the background. A similar test was run with the frame defined by equiluminous colour using a specialized helmet (Figure 1). This gave a strong frame effect. In the second-order motion demonstration below, the red and blue discs are again *always* vertically aligned — notice the green line — but when the line disappears, a large offset may appear with red to the right of blue. 97% of observers saw this offset immediately or with some delay. 3% did not.

**Figure 1.**
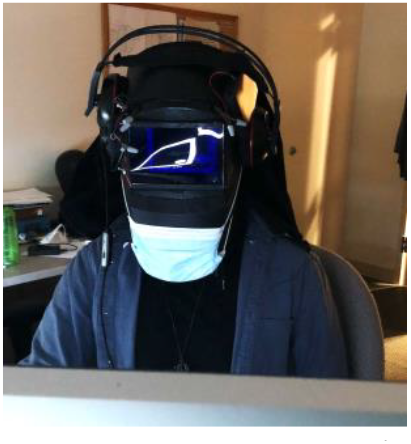
Viewing stimuli with the equiluminizing helmet. The screen display is reflected off the front blue filter. See Connolly et al, 2017.

**Movie 1.1.3.**
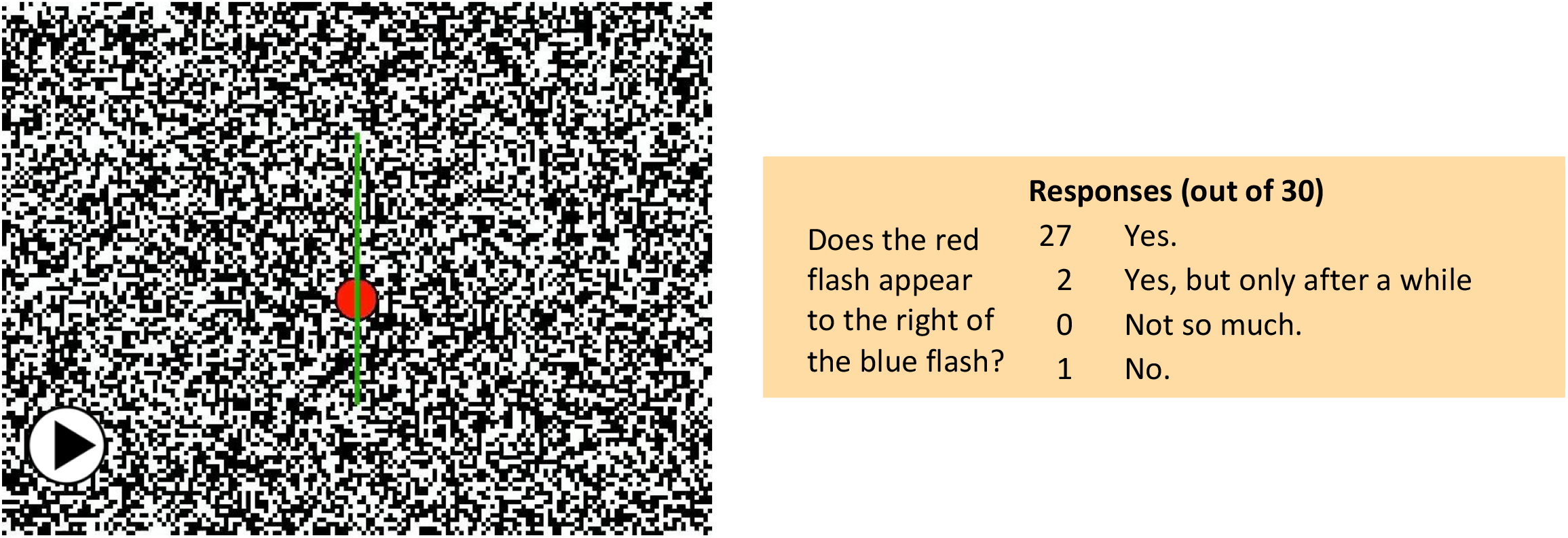
Click to start.

#### 1.1.4. Two frames at once

Here, two frames move opposite directions with all four flashes aligned horizontally. 97% of observers reported the vertical offsets in both frames immediately or with some delay. This shows that the frame effect is not due to eye movement artifacts. Interestingly, smaller effects are seen when comparing positions across the frames.

**Movie 1.1.4.**
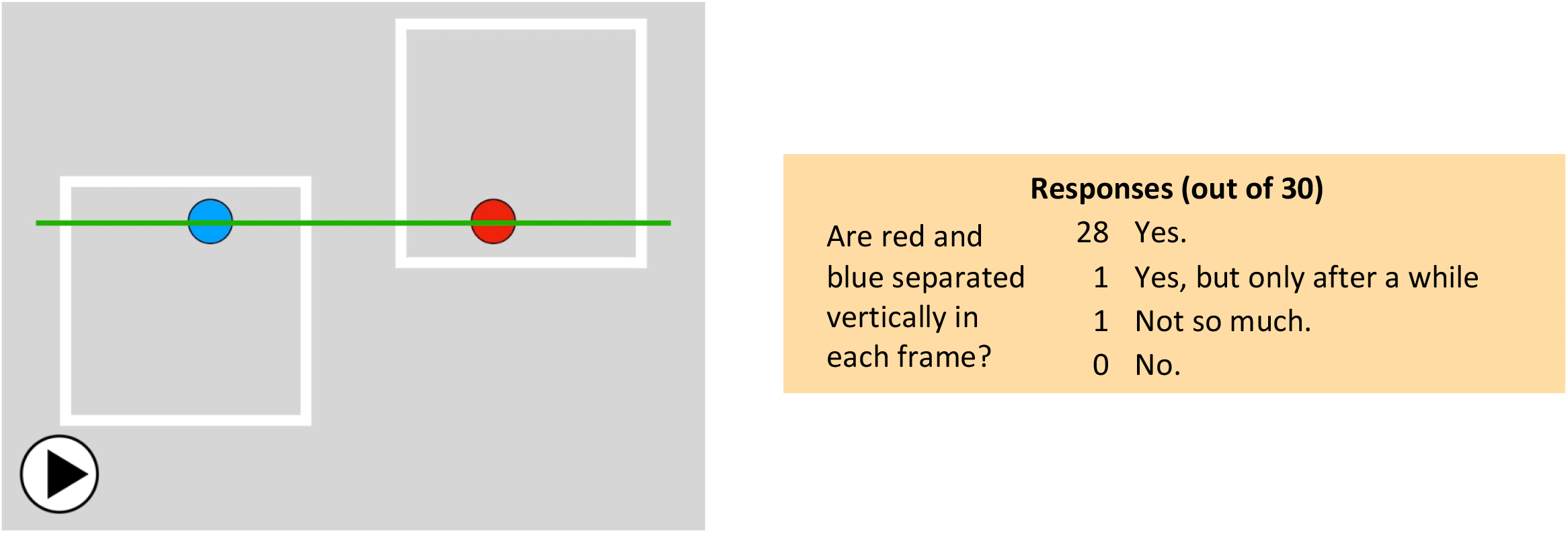
Click to start.

### 1.2. Is the frame effect due to displacement or motion?

We find that both the displacement and the motion generate position offsets, but they act independently. The *displacement* effect is independent of speed and dependent on the size of the displacement (Özkan et al., 2021); whereas, the effect of pure *motion* is speed dependent, independent of path length (Cavanagh, MacLeod, &Anstis, 2021). Pure motion is tested here using reverse apparent motion (Anstis, 1970) and the stimulus offers no discernible landmarks to carry displacement information. The dots move one way but the motion goes the opposite direction. 97% of observers reported that the flashes, which are always vertically aligned, appear displaced in agreement with the perceived direction and not the physical displacement. 3% reported not seeing any shift.

**Movie 1.2.**
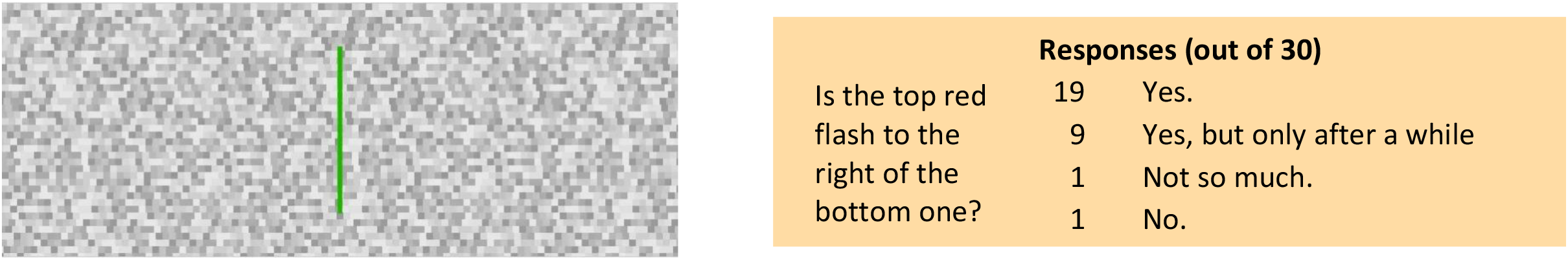
Click to start.

### 1.3. What moving backgrounds can produce the frame effect?

Any moving background appears to be sufficient, from natural scenes, to outline squares and random dots. Even two discs moving in tandem can be effective. The red and blue discs are always vertically aligned as shown at the beginning by the green line, but red may appear shifted to the right of blue once the line disappears. 90% reported the offset immediately or with some delay, 7% did not see it.

**Movie 1.3.**
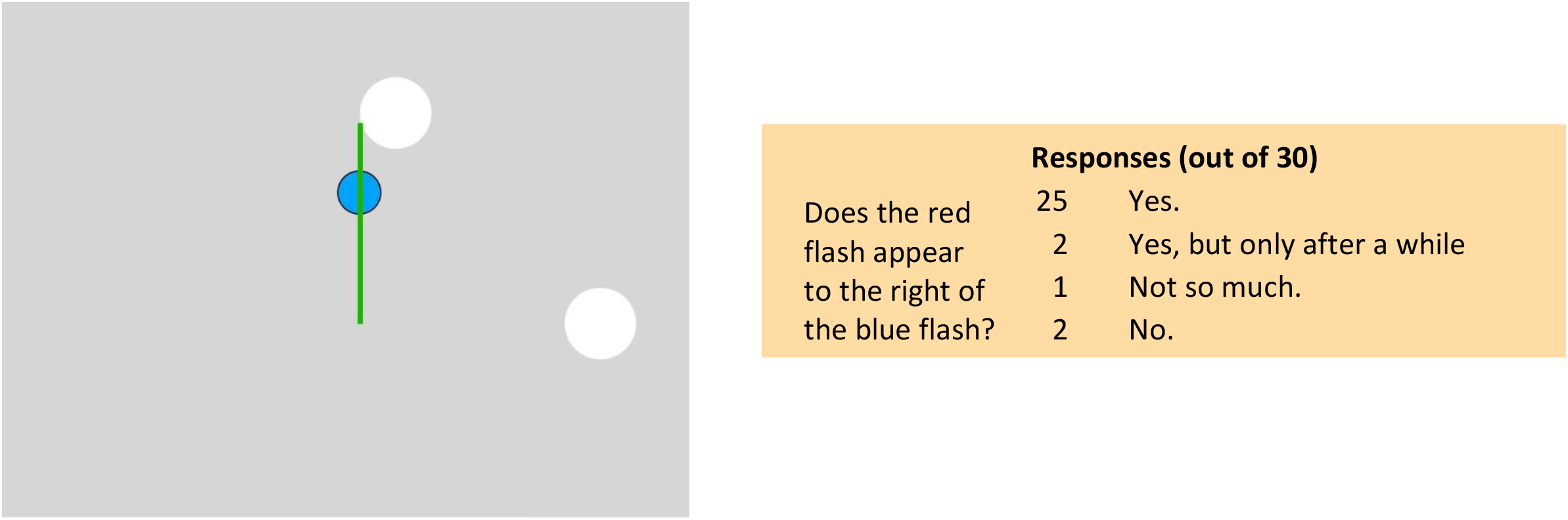
Click to start.

#### 1.4.1. Can the background change while moving?

The background can distort and rotate while still preserving the frame effect. It appears sufficient that some “thing” has displaced. It may change as it displaces. This may be related to the tolerance of apparent motion to shape and feature changes between the first and second position. Here the shape is changing during the motion but once the green line fades, the red disc appears to the right of the blue, even though they are vertically aligned. 93% reported seeing the offset immediately or with some delay, 3% did not see it.

**Movie 1.4.1.**
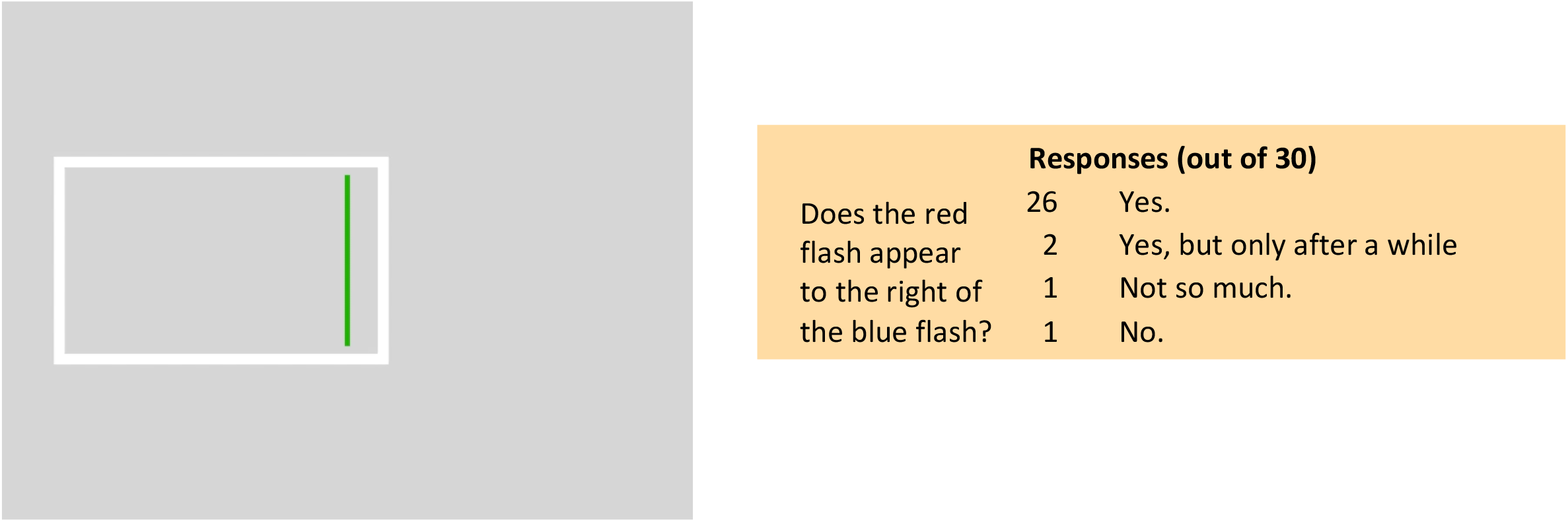
Click to start.

#### 1.4.2. Can the background rotate while moving?

Here the shape rotates while displacing. The effect (red to the right of blue) may be reduced for some observers. 69% reported seen the offset immediately or with some delay, 17% did not see it.

**Movie 1.4.2.**
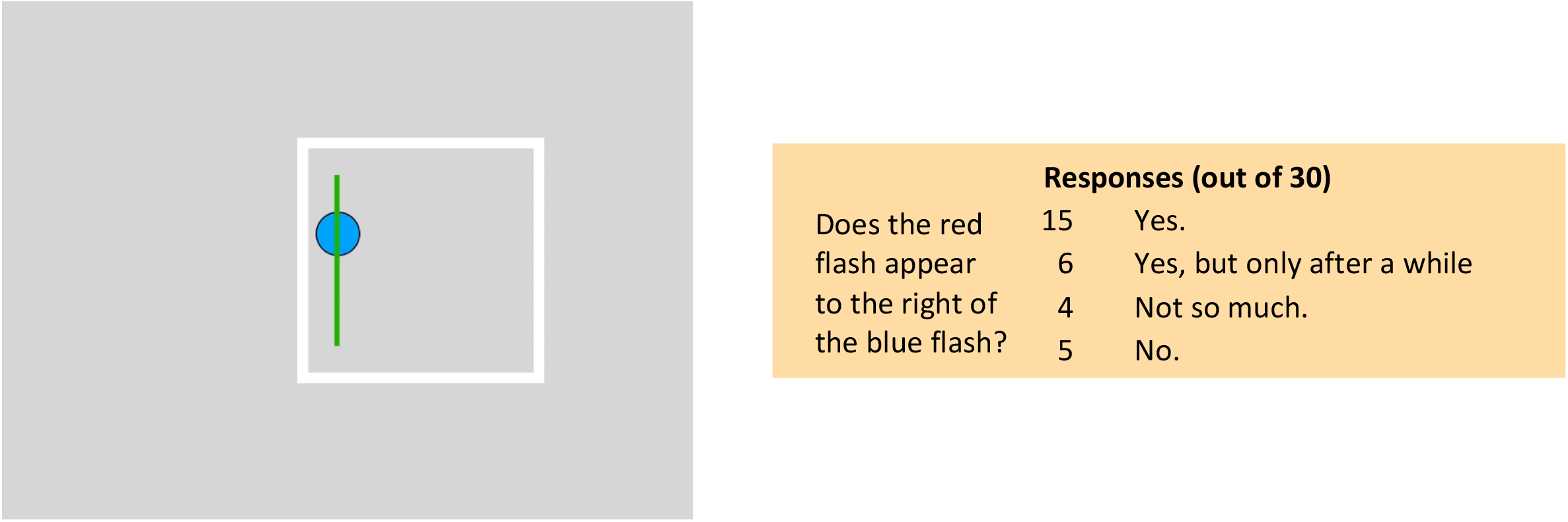
Click to start.

#### 1.5.1. Does the displacement have to be linear?

The backgrounds can follow nonlinear paths while still preserving the frame effect that is driven solely by the displacement between initial and final locations when the two flashes appear. Complicated paths are less effective though. Here is a path that follows a semicircular arc. 86% reported seeing the effect immediately or with some delay, 7% did not see it.

**Movie 1.5.1.**
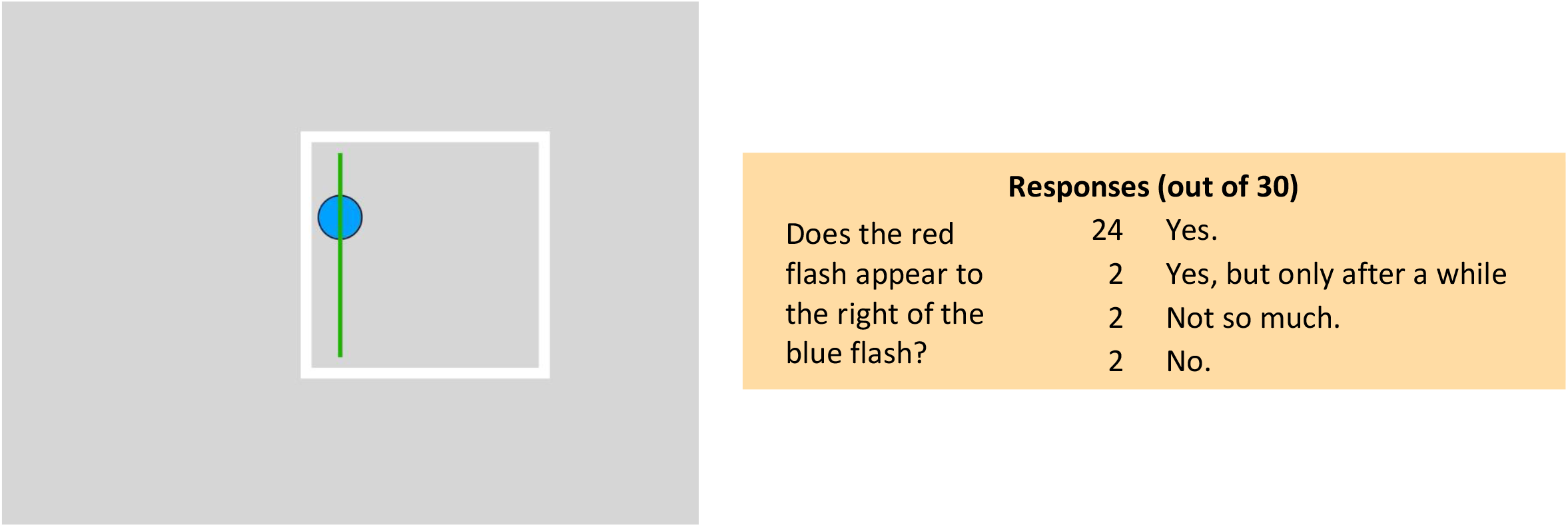
Click to start.

#### 1.5.2. Does the displacement have to be linear?

Does the flashed probe shift because it is pulled by the initial motion of the frame or is its shift a result of the frame’s overall path? We can test this with a nonlinear path that starts out in the direction opposite to the final displacement (Fig. 2) likeNthe one below. The initial motion here is to the right but in the end, the frame has moved to the left. If the shift is caused by the initial motion, the red will appear to the left of blue. If it is caused by the overall displacement, the red will appear to the right of the blue. The outcome was equivocal though, 41% reported seeing red to the right, but 47% reported no offset.

**Figure 2.**
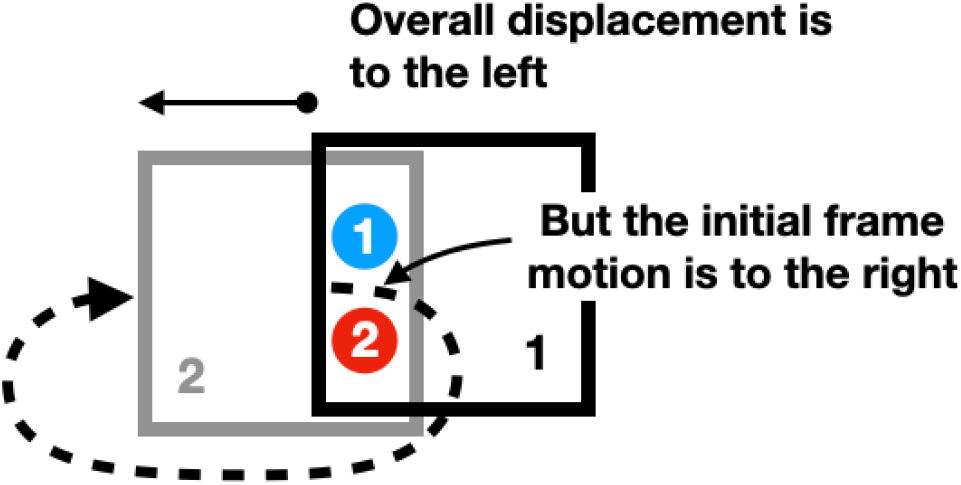
Direction reversing path.

**Movie 1.5.2.**
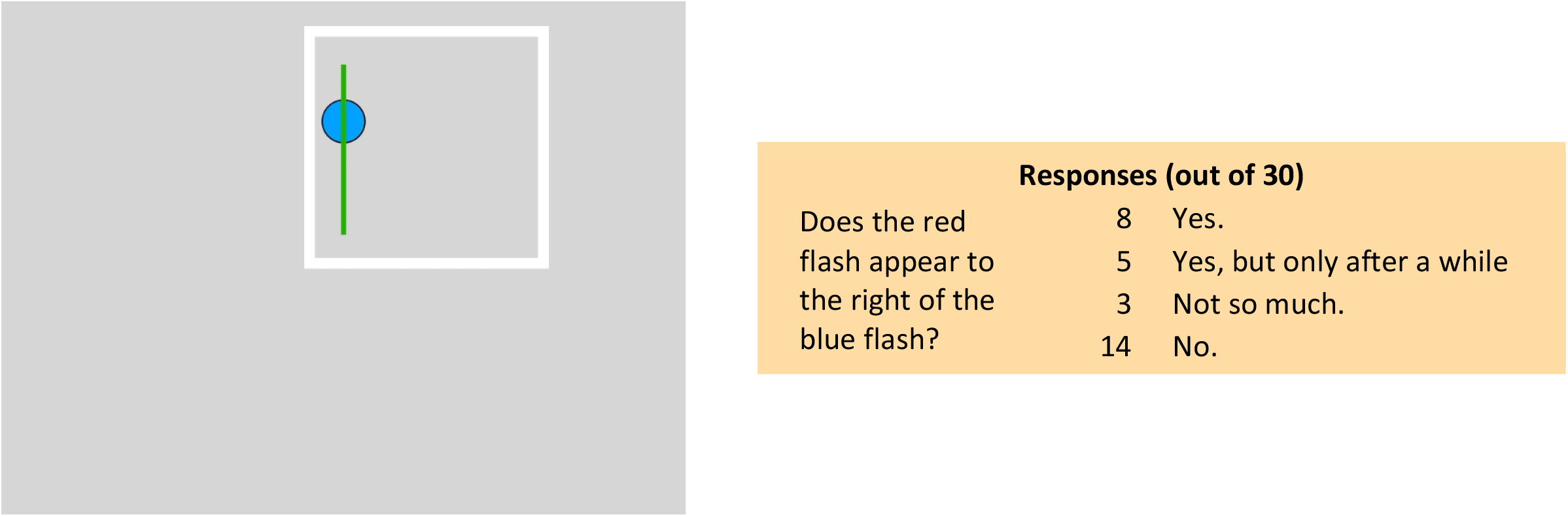
Click to start.

### 1.6. Does the displacement have to have transient reversals?

The backgrounds can follow a continuous circular path with no reversal transient while still preserving the frame effect. 83% saw the offset immediately or after some delay in this movie, 7% did not.

**Movie 1.6.**
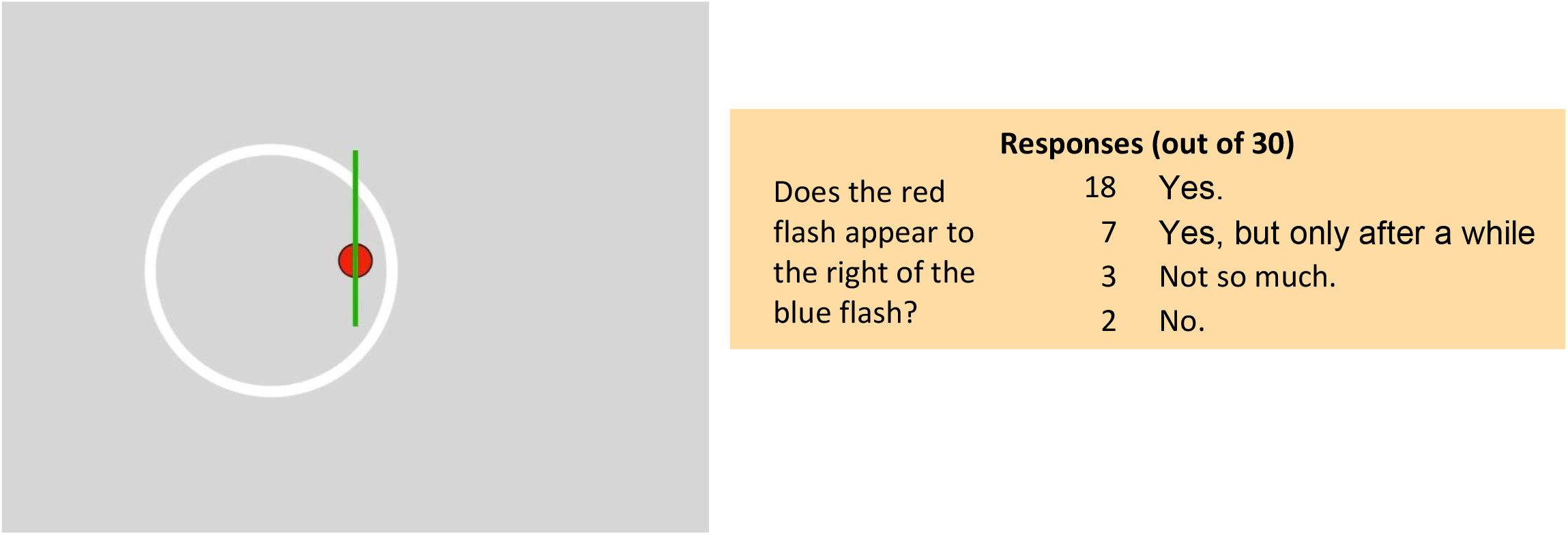
Click to start.

#### 1.7.1. Attention: What happens when there are two frames?

When there are two frames moving in opposite directions, the flashed discs may initially appear vertically aligned, as they are. But **attention** to the light or dark frame can make the red dot appear to the left of the blue (attend to light frame) or to the right (attend to the dark frame). 63% saw the two flash offsets reverse when they switched attention, 23% saw no offset, 13% saw an offset but could not reverse it.

**Movie 1.7.1.**
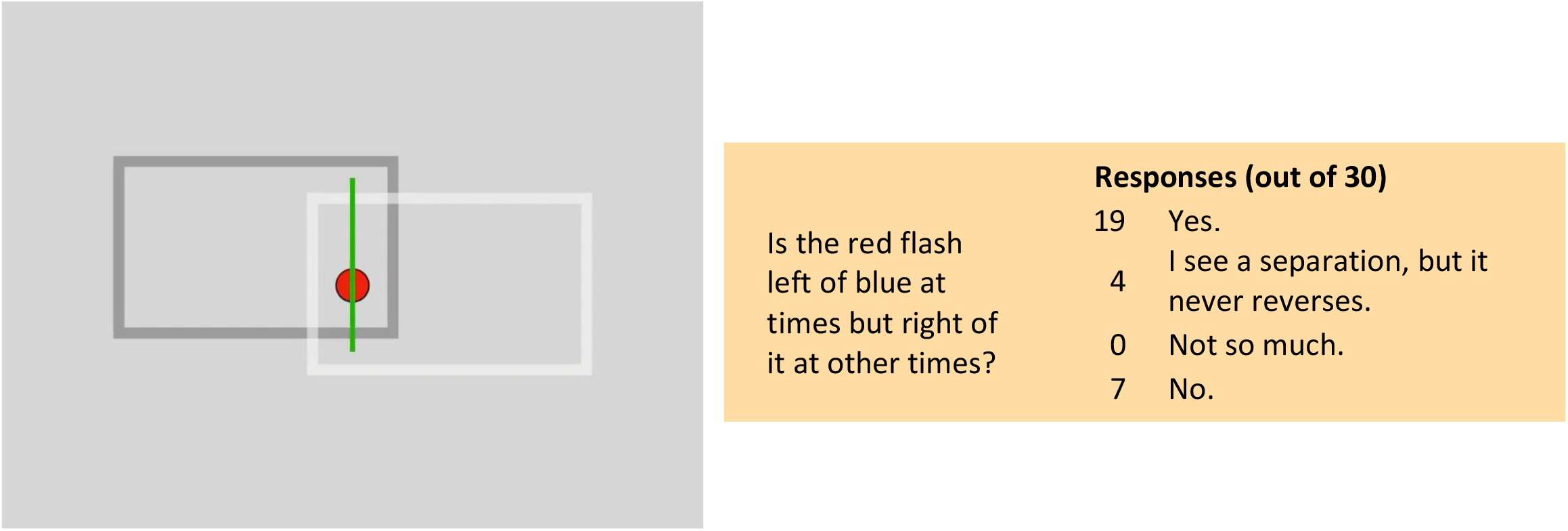
Click to start.

#### 1.7.2. Attention: What happens when there are two frames?

Here there are two frames moving orthogonally, one within the other. The flashed discs may initially appear displaced vertically due to the motion of the inner frame, so blue appears to be above red by more than its physical offset (indicated initially by the green square). The inner frame dominates. However, attention to the outer frame may make the flashed dots appear displaced horizontally as well (blue left of red). This may take some time as the inner frame seems to be more effective — perhaps because it is closer to the flashed dots. 73% were able to see both offsets. 27% saw only one or the other.

**Movie 1.7.2.**
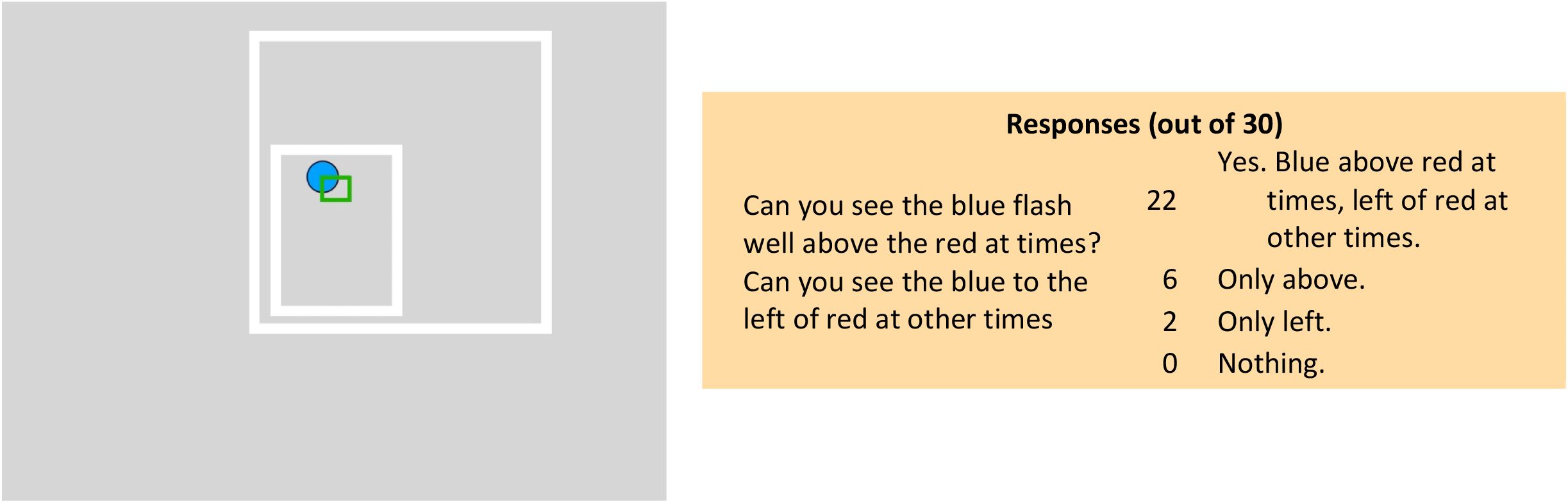
Click to start.

#### 1.8.1. Is the frame effect driven by global or local motion?

Here is the first example to demonstrate that the frame effect is determined by the global, perceived motion not by the local, physical motion. Here, in an angled parallelogram, the perceived motion is up and down even though the local motion of the nearest contours is left and right. When viewing the whole shape, the two flashes shift apart vertically, consistent with the global motion. However, when viewing is restricted to a horizontal aperture, the left-right motion dominates and red and blue may shift apart horizontally. Indeed, all observers saw the offset as vertical before the occluders were present, then horizontal after they appeared.

**Movie 1.8.1.**
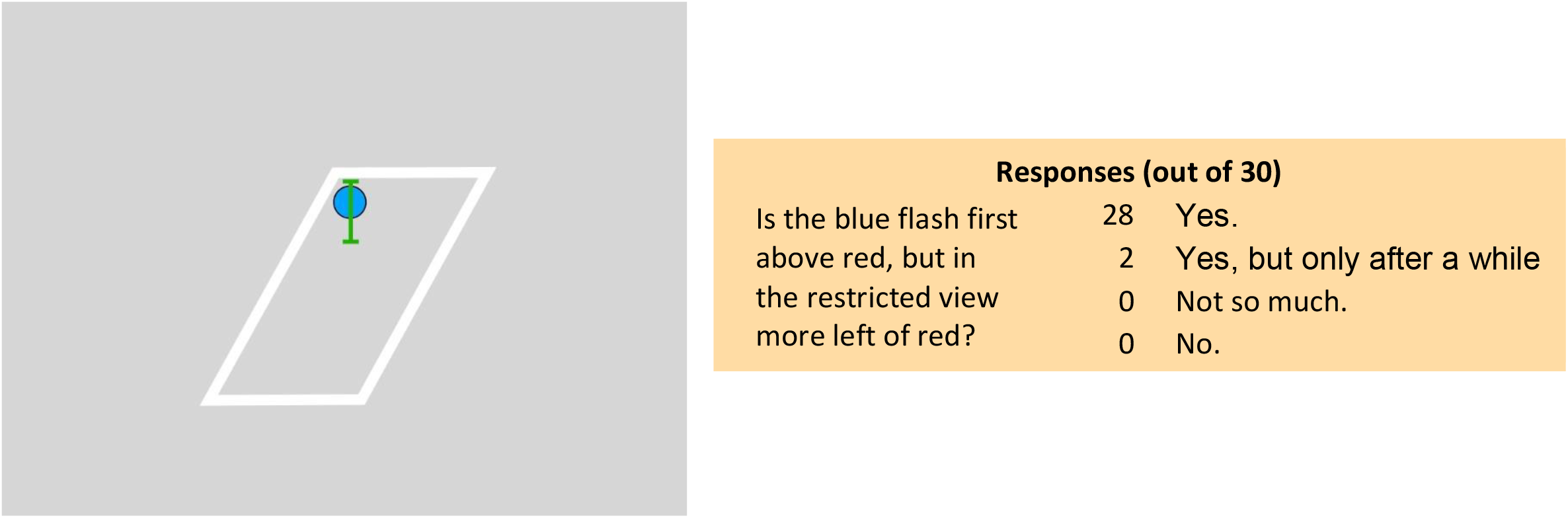
Click to start.

#### 1.8.2. Is the frame effect driven by global or local motion?

In this second example, an occluded diamond shifts left and right (Lorenceau &Shiffrar, VR 1992). With gaze near the flashing discs, the motion of the white contours appears to be up and down and there may be no shift of the two flashes. However, with gaze away to the left or right, a left-right motion may be seen and the two flashes should then shift apart horizontally. 67% of observers did report the offset when looking peripherally, 10% saw no offset no matter where they looked.

**Movie 1.8.2.**
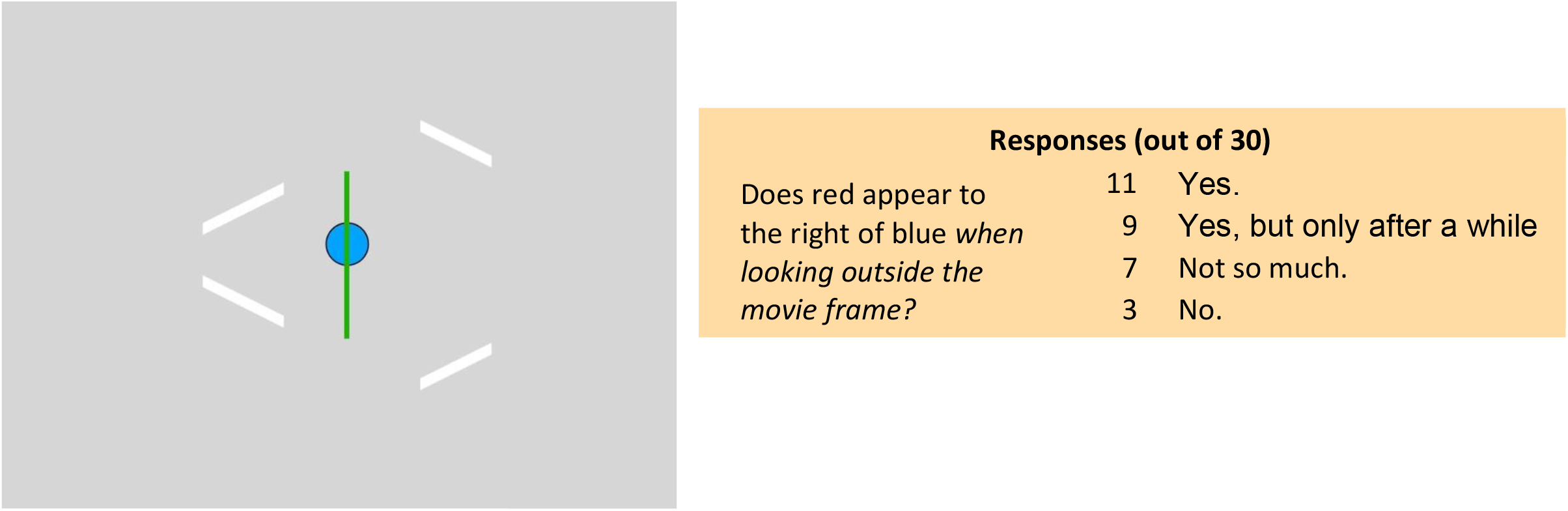
Click to start.

### 2. Where Frames Do Not Work

#### 2.1.1. Frames do not work when there is a nearby spatial anchor

Here there is initially one vertical line. Its presence anchors the locations of the two flashes and 73% of observers report little or no shift indicating that the nearby static anchor has suppressed the effect of the frame, perhaps because it provides a strong position reference. However, if more lines are added, the frame effect returned for 47% of observers, suggesting that the original line lost its position signal as it became an indistinguishable part of a texture. Interestingly, 27% of observers already saw an offset between the two flashes when only one line was present, and they saw the same offset when all the lines were present. 17% saw no offset in either configuration.

**Movie 2.1.1.**
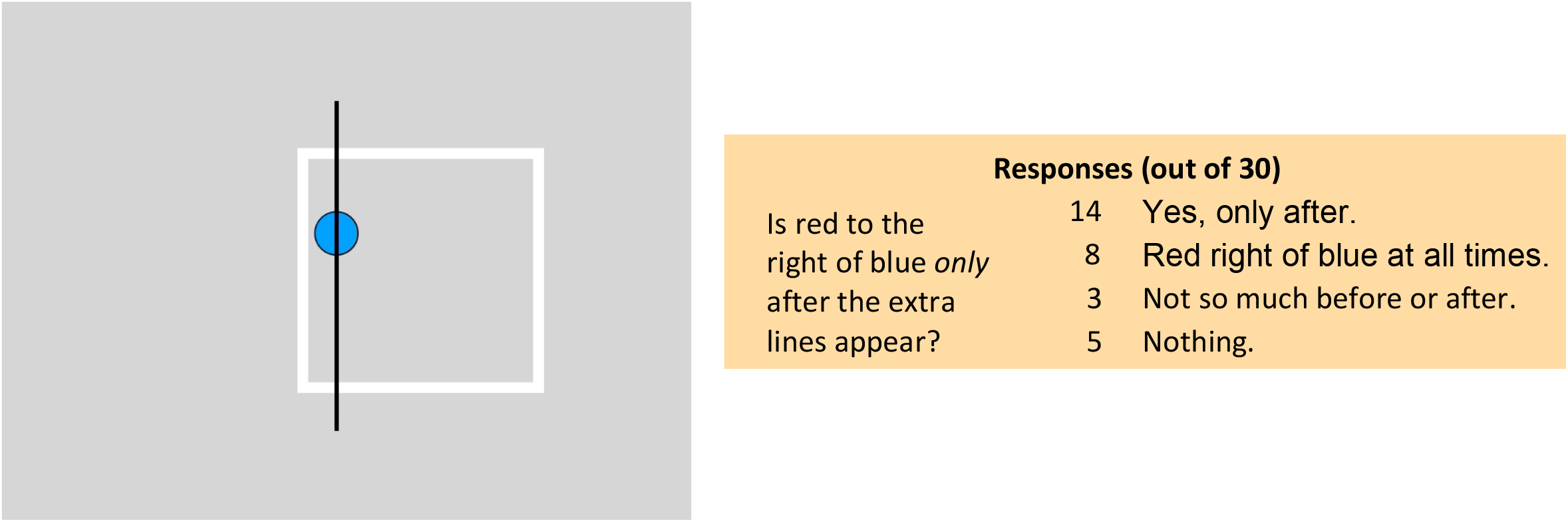
Click to start.

#### 2.1.2. But static textures do not act as spatial anchors

The previous example showed that not all nearby static features will suppress the frame effect, at least if they form a regular texture. Here is a second counterexample where the arrangement of static features is random. The frame effect still works for 90% of the observers despite the nearby stationary spots the texture provides. This shows that the texture does not have to be uniform to lose its position input to the flash judgments. Only 3% of observers reported no offset here.

**Movie 2.1.2.**
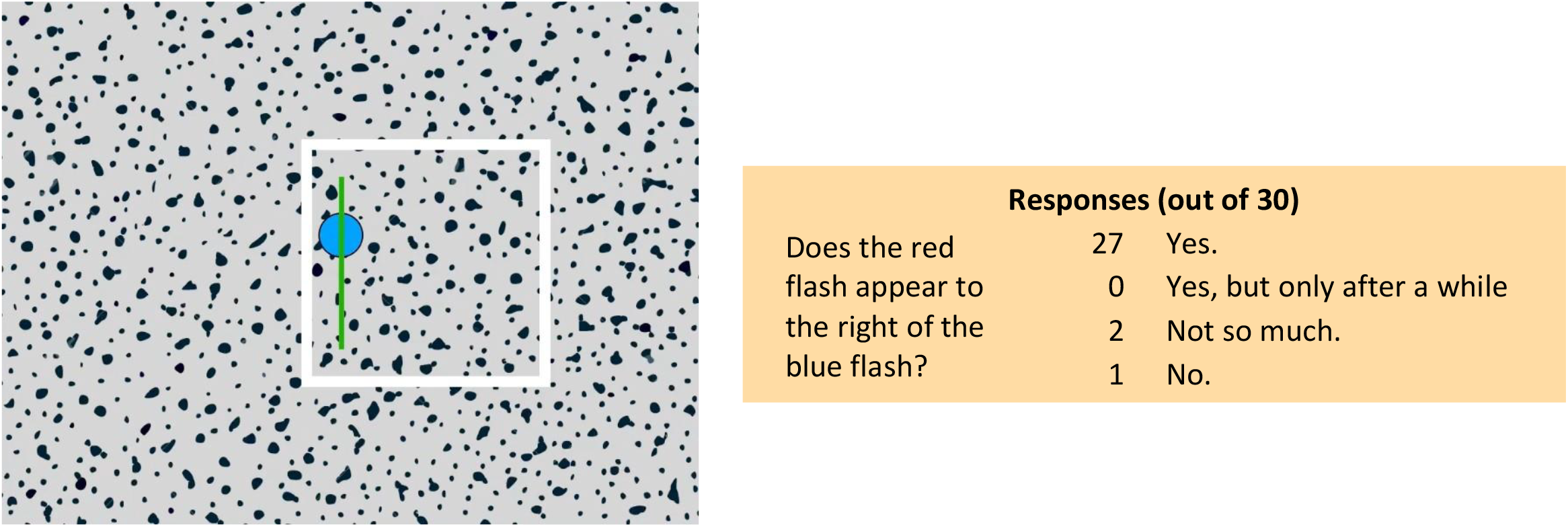
Click to start.

#### 2.2.1. When the frame rotates around the flashes

If the frame rotates around the tests, here by 180°, the frame effect appears to be suppressed for many observers (43%) — red is not seen to the right of blue. However, some observers (28%) did see an offset here. Notice that the start and end positions of the frame are the same as in the original effect. However, the path between the two end points is quite different, with a rotation around the flash locations rather than a translation across them. Possibly, the continuous presence of one edge near the flashes acts as an anchor, at least for some observers.

**Movie 2.2.1.**
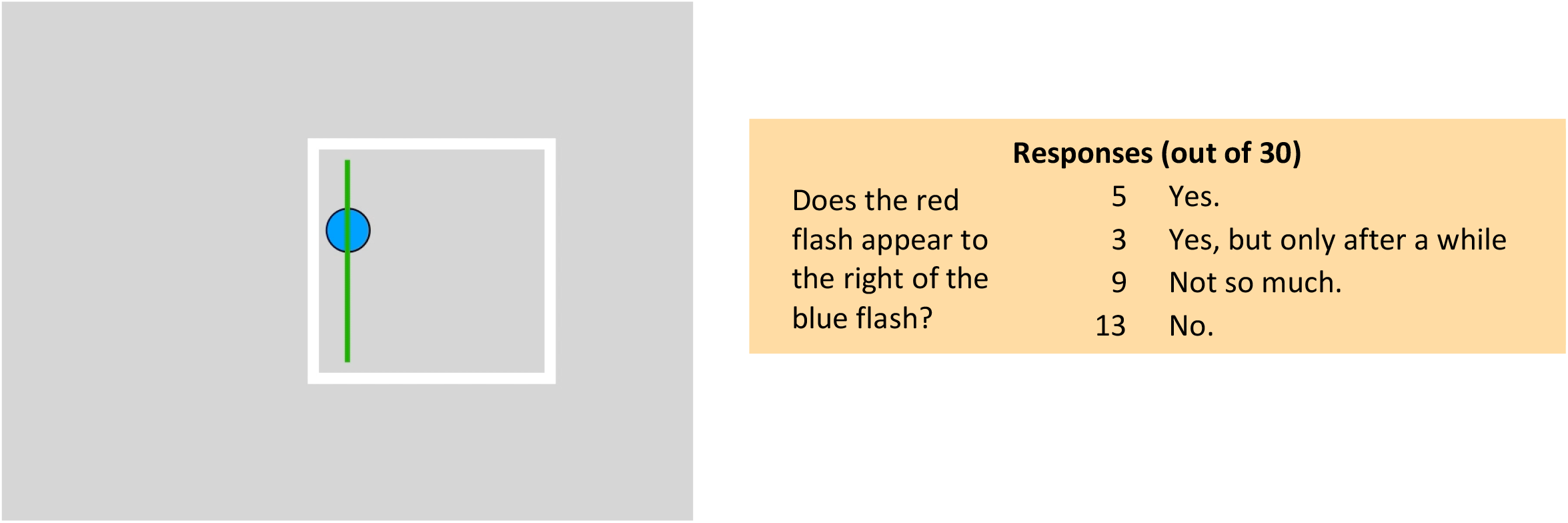
Click to start.

#### 2.2.2. When the frame rotates around the flashes

Here is a similar case but with 270° of rotation. Now, 70% see no offset and the rest, not much. If anything, red may be seen to the left of blue. The start and end position of the frame are again the same as in the original effect, showing that it is not only the end positions that count.

**Movie 2.2.2.**
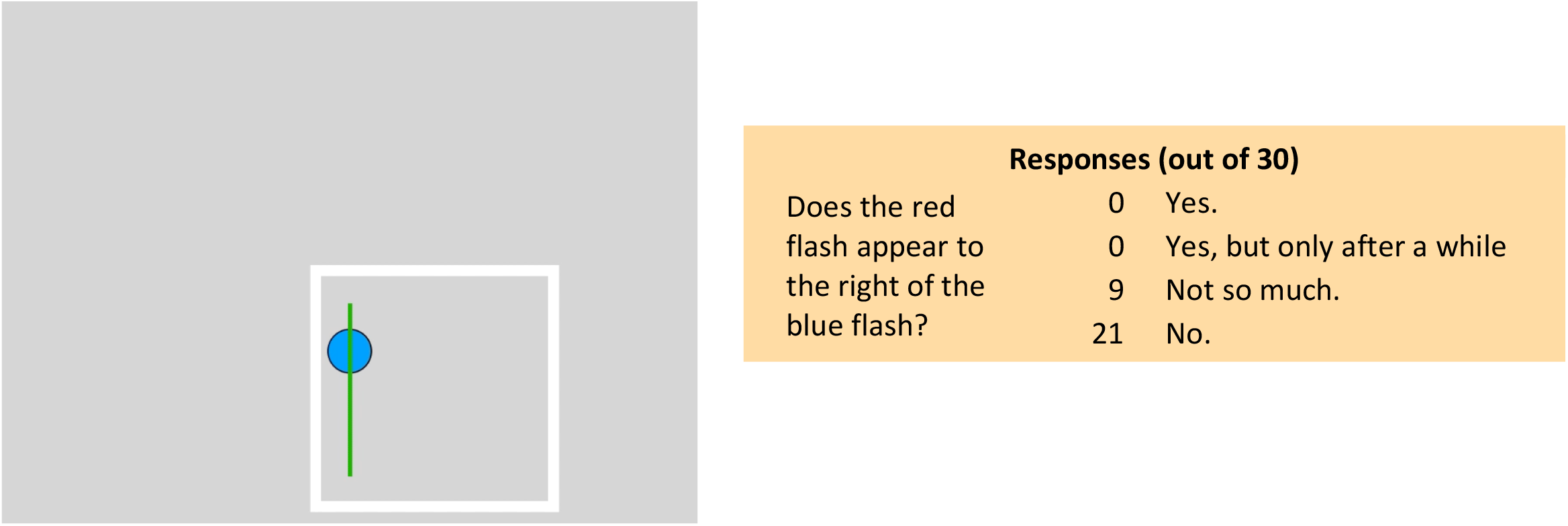
Click to start.

### 2.3. When the frame flips in 3D

Instead of rotation in the plane, the frame now flips out of the plane. Here, 93% of observers report little or no effect, and only 7% see an effect. In fact, you might see red to the left of blue, suggesting that the edge that stays near the flashes serves as a frame or anchor and positions are seen relative to it rather than relative to the flipping frame: red is to the left of that edge when it flashes, blue to its right when it flashes. The start and end position of the flipping frame are the same as in the original translating frame. This shows again that it is not only the end positions that count.

**Movie 2.3.**
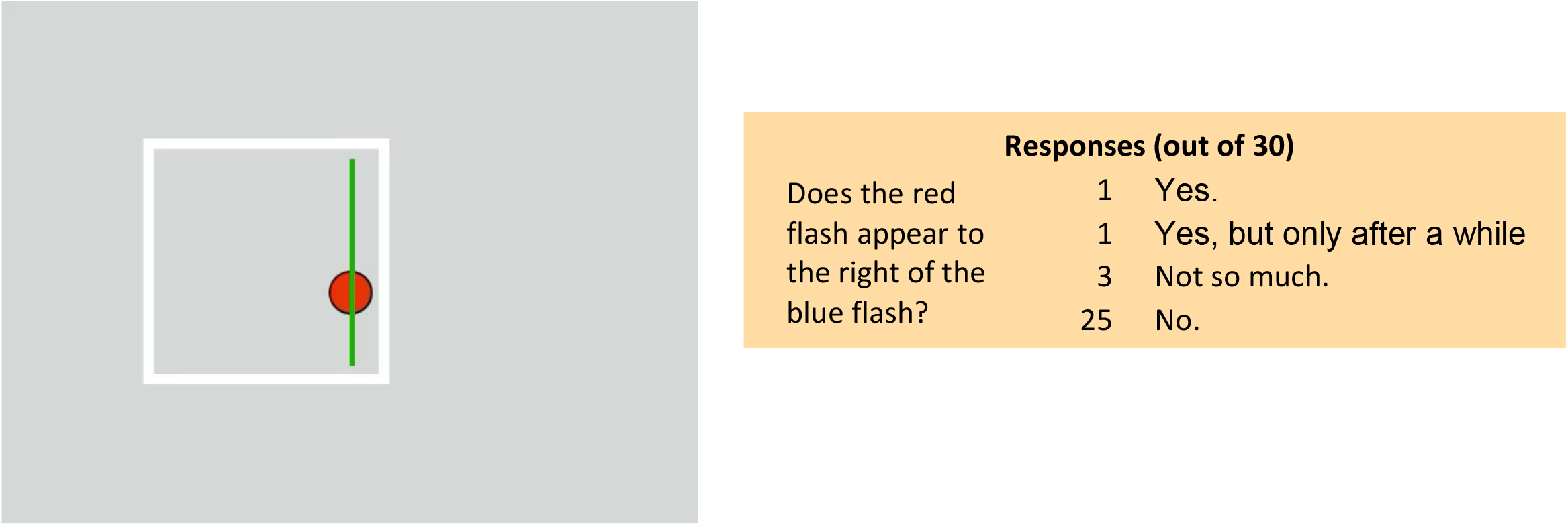
Click to start.

### 2.4. When the probe is continuous

Here the frame moves back and forth in its normal fashion, but when the probes are on continuously (rather than flashed), 93% of observers report little or no visible displacement of the continuous probes. 7% report seeing some shift. The continuous probes alternate with the flashed probes to show the difference. The stationary probes were Duncker’s (1929) original paradigm, although he used a single dot not two. With a continuous probe, the moving frame only produces a small, reversed motion in the probe (induced motion) when the frame’s motion is so slow it is near the motion threshold (e.g., Nakayama &Tyler, 1978).

**Movie 2.4.**
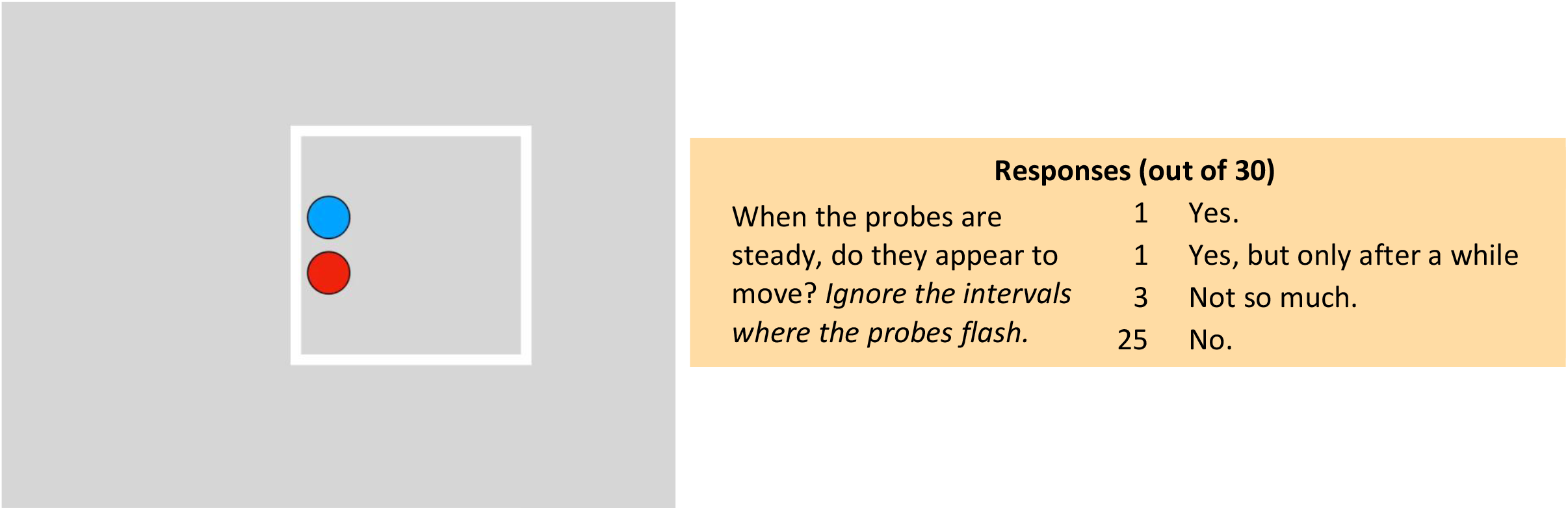
Click to start.

## Conclusions

Previously, the frame effect has been shown to be a remarkably strong illusion, separating aligned flashes by as much as the distance the frame travels (Wong &Mack, 1981; Özkan et al., 2021). Here we explored a wide number of factors to evaluate their influence on the effect. The observations that we collected were only rudimentary subjective reports (e.g., yes, I see it) but they did help identify which stimuli gave strong effects and which gave weak, ambiguous, or no effects. In particular, we found that the frame effect was quite robust and held up for smoothly or abruptly displacing frames and for second-order as well as for first-order motion. It held up even when the frame changed shape or orientation between the endpoints of its travel. The frame’s path could be non-linear, even circular, although the more complex paths were less effective. Two examples using aperture effects (movies 1.8.1, 1.8.2) showed that the separation of the flashed tests was driven by the perceived motion, not the physical motion. Moreover, when there were competing, overlapping frames (movies 1.7.1, 1.7.2), the effect was determined by which frame was attended although not all observers were able to switch attention between the frames.

These results suggest that the frame effect depends on initially tracking the moving frame. We assume that it is acquired and tracked by attention in the same fashion as a target in the multiple object tracking (MOT) task. The result with the overlapping frames supports the role of attention in this tracking – for many observers, the two overlapping frames did not cancel each other’s effect; instead, the offset between the flashed tests depended on which frame was attended. The unattended frame then became ineffective. MOT and the frame effect also share an indifference to shape changes. Radical distortions in shape did not deter the frame effect in movie 1.4.1 and shape changes of targets in MOT do not affecting performance except when the shapes took on the properties of fluids (vanMarle &Scholl, 2003). Apparent motion shows a similar tolerance to shape changes (Kolers &Pomerantz, 1971; Kolers &von Grünau, 1976; Zhou et al., 2003). In all these cases, it is a persisting “object” that is being tracked, one whose continuity depends primarily on spatiotemporal proximity rather than fixed shape. Kahneman, Treisman, and Gibbs (1992) addressed this same point with their concept of an “object file” as a transient representation of an entity that may change its properties over time but remains the same thing. In the case of the frame effect, this entity, the moving frame, is also serving as a reference for the location of events that take place in its neighborhood. Interestingly, this suggests that the frame effect could be a useful tool for evaluating the principle of continuity – offering an objective test based degree of illusory shift of the flashed probes. The frame effect also holds promise for understanding mechanisms underlying visual stability where the displacement of the entire visual scene may acts as a moving frame that stabilizes position as the eyes move.

We also found that there were a number of constraints that limited the effect. An isolated, static anchor near the flashes suppressed the effect but an extended static texture did not. When the frame’s path kept one edge of the frame near the flash locations, the effect was reduced or eliminated. If the probes were continuous (as in Dunker, 1929) rather than flashed, the effect was abolished as well. Indeed, previous studies have shown that a moving frame influences a steady probe only when the frame’s motion is very slow, much nearer motion threshold than the speeds used here (Nakayama &Tyler, 1978; Reinhardt-Rutland, 1988). Possibly, any visible relative motion between the probes and frame shows that the probes do not belong to the frame (do not group with the frame) and so suppresses the frame effect. Alternatively, the continuous unmoving probe may group with the steady background beyond the moving frame, suppressing any effect of the frame.

The earlier article (Özkan et al., 2021) already established that the frame’s motion can separate the perceived positions of the flashes by as much as the frame’s travel — equivalent to being seen in frame coordinates (the locations in the frame where the probes were when they flashed) with the frame stationary. In this case, its motion would be completely discounted. But intriguingly, the frame is still seen to move quite well, albeit over a shortened path. Combined with these earlier results, our new observations lead us to propose a rough organization for the frame effect, laid out in Figure 3, where the frame’s motion acts along two different pathways once the frame has been acquired and its motion tracked.

**Figure 3.**
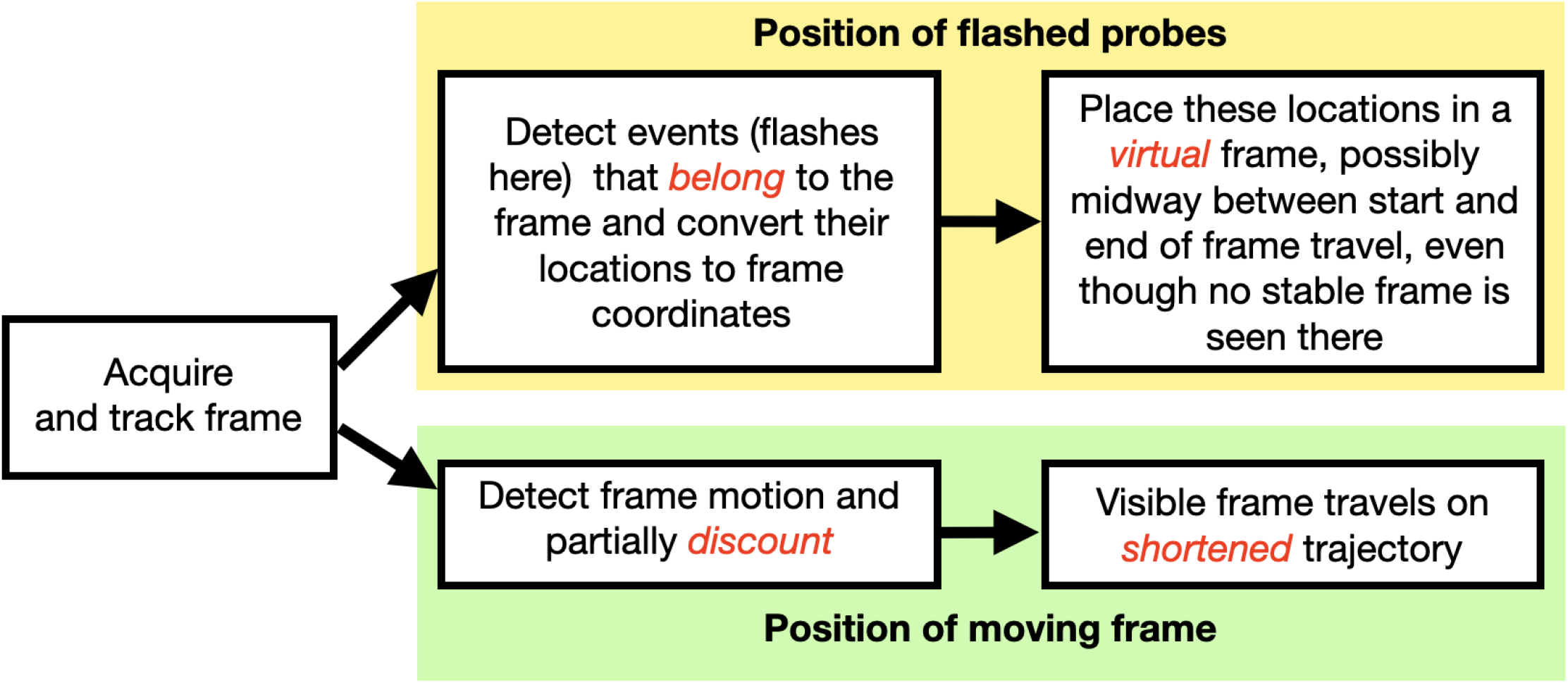
Proposed organization of the frame effect

In the first, the upper path in Figure 3, the frame’s motion acts on the positions of the flashes, contingent on the flashes belonging to or binding with the frame. For example, when attention switched from one frame to the other in movies 1.71 and 1.72, the direction of the separation of the flashes switched as well, indicating that the flashes were seen in the context of, or belonging to, the attended frame. Importantly, the large separations between the flashes matched their separations in frame coordinates – the locations the flashes had relative to the frame when they flashed. Moreover, this illusory separation was seen centered on the frame’s path — as if the flashes were fixed in those locations relative to a static frame that was located in the middle of the path. However, this static frame was clearly not visible. We can think of this unseen frame as a virtual or amodal frame although we have no evidence yet of its existence.

The second pathway, the lower one in Figure 3, concerns the perception of the frame’s motion. The frame does not come to a standstill as it should to be consistent with large flash separations, but its motion is nevertheless attenuated. This “path shortening” has been reported for bounded motion trajectories before (Whitney, Murakami, &Cavanagh, 2000; Sinico, Anstis, &Casco, 2009; Cavanagh &Anstis, 2013) and is apparent here again.

The observations here and the rough outline of how frames work in Figure 3 suggest several new directions for understanding the frame effect. These build on the nearly 100 years of research on the effect of frames on visual perception but bring new questions. How is the visual scene decomposed into hierarchical sets of dynamic frameworks? What are the limits to the changes of the frame that still maintain its continuity? What properties from each frame are inherited by the elements belonging to the frame? What determines when a test flash “belongs” to the frame? The remarkable strength of the frame effect on perceived location also suggests that it can serve as a “workhorse” tool, a visual equivalent to the Stoop task, that can be used to examine the details of how vision encodes the dynamic scenes unfolding in front of us.

## Acknowledgments

NSERC Canada (PC,BMtH), Dartmouth PBS (SS, MM), ANR (MW), UCSD Psychology (SA).

## Notes

https://cavlab.net/Demos/FrameEffect

https://cavlab.net/Demos/FrameExperiment

### Competing Interest Statement

The authors have declared no competing interest.

### Summary of Updates

The introduction, methods and conclusions were expanded and the number of participants in the dataset increased to 30 from 26.

